# Characterization of gustatory receptor 7 in the brown planthopper reveals functional versatility

**DOI:** 10.1101/2020.09.05.284208

**Authors:** Abhishek Ojha, Wenqing Zhang

## Abstract

Insect pests consume tastants as their necessary energy and nutrient sources. Gustatory receptors play important roles in insect life and can form within an extremely complicated regulatory network. However, there are still many gustatory genes that have a significant impact on insect physiology, but their functional mechanism is still unknown. Here, we purified and characterized a gustatory receptor (protein) coding gene, *NlGr7*, from the brown planthopper (BPH) *Nilaparvata lugens*, which is an important insect pest of rice. Our results revealed that *Nl*Gr7 has an active association with various ligands, such as lectins, lipids (phospho- and sphingolipid) and copper. The mass-spectrometry result showed that *Nl*Gr7 is a sugar receptor, and *Nl*Gr7 is validated by different types of insoluble polysaccharides and a varied range of tastants. Furthermore, we observed that *Nl*Gr7-bound ATP hydrolysed on the ATPase activity assay, which indicated that *Nl*Gr7 may be associated with important biological functions in the BPH. The important *Nl*Gr7 for chemoreception has now been characterized in the BPH. We showed that *Nl*Gr7 in the BPH is required for various protein-ligands, as well as protein-sugars interactions, to play crucial roles in this pest. This study will provide valuable information for further functional studies of chemoreception mechanisms in this important agricultural pest.

## Introduction

Arthropods’ gustatory receptors (Grs) are active in leading insect feeding behaviours and in establishing the platform between insect gustation and the environment. The Grs are closely associated with insect olfactory receptors (Ors) (**Robertson et al., 2003**). Grs have been investigated as divergent domains of seven-TM (transmembrane) spanning proteins, which is a typical example of G protein-coupled receptors (GPCRs), but they are noticeably diverse and impart no sequence resemblance with Grs of mammals (**Clyne et al., 2000)**. In addition, the topology of Gr’ seven-TM helices are reversed in comparison to that of the GPCRs (**Zhang et al., 2011**). Although having well-established information on olfaction (**Benton et al., 2006; Robertson et al., 2006**) and increasing documentation of insect Grs, little known about the physiological mechanisms that include understanding of the gustation system.

The *Gr* genes were first established in *Drosophila* (**Clyne et al., 2000**), which is a polyphagous dipteran insect, and forty-three putative *Gr* genes have been revealed in this insect (**Amrein and Throne, 2005**). Grs have been documented into various clades and are named as sugar (**Slone et al., 2007)**, bitter (**Wanner and Robertson, 2008**; **Lee et al., 2009**), carbon dioxide (CO_2_) (**Xu and Anderson, 2015**), GR43a-like (**Kui et al., 2018**), sex attractant (**Shankar et al., 2015**), and unknown (**Kui et al., 2018**) receptors, based on their sequence homologies with receptors that have been discovered in *Drosophila* (**Robertson et al., 2003**) or on an important molecule to which they reacted (**Kui et al., 2018**). Seventy-six and 79 *Gr* genes have been respectively identified in *Anopheles gambiae* (**Hill et al., 2002**) and *Aedes aegypti* (**Kent et al., 2008**), which is a hematophagous dipteran insect. Sixty-five and 197 *Gr* genes have been noted in the lepidopteran insects *Bombyx mori* (**Wanner and Robertson, 2008**) and *Helicoverpa armigera* (**Xu et al., 2016**), respectively. The highest number of *Gr* genes, 220, has been observed in *Tribolium castaneum*, which is a Coleopteran insect (**Richards et al., 2008**). Currently, the majority of research consideration has been focused on the *Gr* genes of *Drosophila* (**Slone et al., 2007; Dunipace et al., 2001; Gardiner et al., 2008; Weiss et al., 2011; Dahanukar et al., 2001; Dahanukar et al., 2007**); however, with fast-growing genome examinations on an additional species of insects (**Robertson and Wanner, 2006; Wanner and Robertson, 2008; Richards et al., 2008; Smadja et al., 2009; Xue et al., 2014**), the investigation is approaching a different spectrum of species. However, there is a need to shed light on the *Gr* genes in an important agricultural insect, such as the brown planthopper. The brown planthopper (BPH), *Nilaparvata lugens* (Stål) (Hemiptera: Delphacidae), is one of the most serious agricultural planthopper species and feeds on rice plants (**Ojha and Zhang, 2019**). Recently, thirty-two putative *Gr* genes in this insect have been reported from our research group (**Kui et al., 2018**). However, the gustatory perception of these *Gr* genes, or the protein macromolecular platform, in the physiology of this insect has remained largely unknown.

In the present investigation, the functional properties of BPH Gr7 (*Nl*Gr7, protein) have been revealed after homologous expression in *E. coli*. Purified *Nl*Gr7 was examined for its association with lectins, lipids, polysaccharides, metals, and ATPase activity assays. These results are essential for understanding the molecular level of taste regulation within the BPH feeding behaviour and improve our understanding of the insect sugar-receptor family, insect-plant interaction and adaptation.

## Materials and Methods

### Insect sample

The BPH strain was maintained and reared on susceptible rice (Huang Hua Zhan) plants as described by Kui et al (2018).

### Bioinformatics

The *Nl*Gr7 (protein) sequence was examined for different parameters, including the molecular weight, theoretical pI, amino acid composition, atomic composition, extinction coefficient, estimated half-life, instability index, aliphatic index and grand average of hydropathicity, by using an ExPASY ProtParam tool (https://web.expasy.org/protparam/). The TOPCONS tool was used to predict a secretory signal sequence and the transmembrane helix in the *Nl*Gr7 (http://topcons.cbr.su.se/). Furthermore, the *Nl*Gr7 sequence was searched with other known annotated insect Grs for sequence identity and to build a phylogenetic tree using protein BLAST tools from the NCBI weblink (https://www.ncbi.nlm.nih.gov/blast/treeview/treeView).

### Cloning, expression, purification and biotin labelling of *Nl*Gr7

The total RNA of the female adult BPH was isolated by using a TRIzol reagent (Invitrogen, Carlsbad, CA), and one μg RNA was converted to cDNA by using the PrimeScript™ RT-PCR Kit (Takara, Japan) following the manufacturer’s instructions. Then, 10 ng cDNA was used as a template for PCR to amplify *NlGr*7 using the forward primer 5’-CATATC-ATGTTGTTACTGACAATTGTTTTGATT-3’ containing an *NdeI* sequence and the reverse primer 5’-GAATTC-TCACTGTGTTCTGCGCGTTT-3’ containing an *EcoRI* sequence. A PCR was carried out in 25 μl containing 200 μM dNTPs, 5 U *Taq* DNA polymerase (New England Biolabs, NEB, Inc, USA) and 13 μM of each of the primers. The PCR conditions were 95°C for 2 min, followed by 35 cycles at 95°C for 30 s, 55°C for 30 s, and 72°C for 30 s, with a final extension step at 72°C for 5 min. Amplicon (∼730 bp) was analysed on 1% (w/v) agarose gel, was gel-purified using HiPure Gel Pure DNA Micro kit (Magen, China) following the manufacturer’s instructions and was quantified using the NanoVue Plus spectrophotometer. The PCR product was digested with *NdeI* and *EcoRI* and was ligated to pMAL-c5X (NEB, Inc, USA). *Escherichia coli* (*E. coli*, DH5α) was used for transformation of the pMAL-c5X-*NlGr*7 plasmid. The transformed bacteria were selected by screening the colonies on ampicillin (50 μg/ml) containing media and plasmid purification. Then, the colonies were further analysed by restriction enzyme digestion and PCR. The *NlGr*7 gene of the recombinant plasmid was sequenced by the Sanger method. The expression host *E. coli* BL21(DE3)pLysS was used as a transformation host for the pMAL-c5X-*NlGr*7 vector. A single colony of transformed *E. coli* BL21(DE3)pLysS with pMAL-c5X-*NlGr*7 was incubated overnight in a shaking incubator in 500 ml Luria Bertani (LB) medium containing ampicillin (50 μg/ml) and chloramphenicol (34 μg/ml) at 37°C with constant agitation (200 rpm). The transformed cells at 25°C (OD_600_ = 0.6) were induced with 0.5 mM isopropyl b-D-thiogalactopyranoside (IPTG; Sigma, USA), which resulted in the expression of the maltose-binding protein (MBP)-tagged *Nl*Gr7 protein. After 8 h of induction, the cells were harvested at 5,0009xg for 15 min. The pellet was resuspended in a lysis buffer containing 50 mM Tris– HCl, pH 8.0, 200 mM NaCl, 3 mM β-ME, 10% v/v glycerol, and 0.15 mg/ml lysozyme. Cells were lysed by sonication (50 Hz, 20 cycles, with each cycle consisting of 20 s on- and 120 s off-time, 4°C), and the sonicated suspension was centrifuged at 20,0009xg for 30 min. The cleared supernatant was applied to amylose beads (NEB, Inc, USA), and protein was eluted with buffer 25 mM Tris–HCl pH 8.0, 200 mM NaCl, 3 mM β-ME and 10 mM maltose. Eluted *Nl*Gr7 protein fractions were checked by SDS-PAGE (12%), and the pure fractions were pooled. The eluted *Nl*Gr7 protein fractions were applied on a 10 kDa cut-off Centricon centrifugal device (Merck Pte. Ltd., Germany) for desalting against buffer (25 mM Tris–HCl, pH 8.0, 10 mM NaCl) and were concentrated to 56.75 mg/ml.

Cleavage of the MBP tag from *Nl*Gr7 was completed according to manual instructions of the pMAL protein fusion and purification system (NEB, Inc, USA). The tag (MBP) was removed by incubating with Factor Xa at 4°C for 3 h. The cleaved *Nl*Gr7 protein was concentrated by using a 10 kDa cut-off Centricon centrifugal device (Merck Pte. Ltd., Germany) and was purified by affinity chromatography on amylose resin (NEB, Inc, USA) equilibrated with 20 mM Tris-HCl, pH 8.0, 200 mM NaCl, and 2 mM DTT; then, it was purified by a benzamidine column (GE Healthcare, USA) equilibrated with 20 mM Tris-HCl, pH 8.0, 200 mM NaCl and 2 mM DTT. Unbound *Nl*Gr7 fractions were checked by SDS-PAGE and were concentrated to 10.00 mg/ml and stored at −80°C. Subsequently, the purified protein of *Nl*Gr7 was analysed using LC-MS/MS (Q-TOF). The labelling of *Nl*Gr7 with biotin was completed as described by Ojha et al (2014). The expression and purification of the MBP (as a control protein) was completed as described above. The protein was concentrated to 61.77 mg/ml and stored at −80°C.

### Protein-lectin-binding assay

To study lectin binding with *Nl*Gr7, 96-well Nunc microtiter plates (Fisher Scientific, Waltham, MA) were coated with *Nl*Gr7 in (100 ng; 50 µl/well) overnight (15-16 h, at 4°C). The plates were washed to remove unbound proteins, were blocked with 3% bovine albumin serum in Tris buffer and were incubated for 2 h at 100 rpm at room temperature. Additional biotinylated lectins (20 µg/ 50 µl; Vector Labs, Burlingame, CA) were added to each well and were incubated for 2 h at room temperature. Then, the plates were washed thrice with Tris buffer containing 0.05% Tween-20 to remove unbound or weakly bound biotinylated lectins, followed by incubation with streptavidin-HRP (1:2,000, 1 h, room temperature). The HRP activity was measured using a 50 μl TMB substrate solution (TransGen Biotech Co., LTD, China) containing 0.06% of 30% v/v hydrogen peroxide (H_2_O_2_) in each well. After 20 min, the reaction was stopped by adding 50 μl sulfuric acid (0.5 N) to each well. The absorbance was measured at 450 nm using an ELISA plate reader. The appropriate negative control was included, and the tests were run in quadruplicate.

### Protein lipid overlay (PLO) assay

To test the binding ability of MBP-*Nl*Gr7 and MBP to various lipids, the well-established protein-lipid overlay (PLO) assay was performed using PIP and sphingo strips (Molecular Probes, USA) according to the manufacturer’s instructions. These strips contained 100 pmol of various phospholipids and sphingolipids, which were spotted and immobilized on a nitrocellulose membrane. The purified MBP-*Nl*Gr7 was overlaid on PIP and sphingo strips. The purified MBP was used as an experimental control. Approximately 25 µg/ml protein was incubated with the strips in TBS-T (contained 3% BSA, Biotechnology Co. Ltd, Guangzhou, Xiang Bo) overnight at 4°C. The strips were then washed with TBS-T/BSA three times with gentle agitation for 10 min each wash, at room temperature. The MBP-*Nl*Gr7 and MBP interaction with spotted lipids were detected by subsequently blocking the strips in TBS-T buffer (10 mM Tris, pH 7.5, 70 mM NaCl, and 0.1% Tween) with 3% BSA and then incubating the strips in 1:2000 dilution of monoclonal anti-MPB-HRP antibody (NEB, Inc, USA) in a blocking buffer for 1 h at room temperature. After thorough washing, a horse reddish peroxidase signal was detected by using 3,3’-diaminobenzidine tetrahydrochloride (DAB, Thermo Scientific) to yield an insoluble brown product. The intensity of the signals was analysed by using ImageJ.

### An electrophoretic mobility shift assay (EMSA) for a metal-binding assay

The metal-binding properties of *Nl*Gr7 were confirmed by mixing purified *Nl*Gr7 (approximately 0.75 μg/5 μl) with equal volumes of the following solutions: 0.5 mM (final concentration) EDTA or 0.015, 0.05, 0.15, 0.5, or 1.0 mM metals (CuSO_4_, and CaCl_2_). Each mixture was incubated for 30 min at 25°C, was added to 10 μl Laemmli sample buffer, and then was subjected to 12% SDS-PAGE under reducing/non-heating conditions. As a negative control, we used the maltose-binding protein for metal-binding assay.

### ATPase activity assay with purified *Nl*Gr7

The ATPase activity assay was performed in 96-well microtitre plates by using an ATPase/GTPase activity assay kit (Sigma-Aldrich, USA) according to the manufacturer’s instructions. An aliquot of *Nl*Gr7-purified protein (6.25, 12.5, 25, 50, 100 μg/well) was mixed with a 5 μl assay buffer to make 10 μl of the ATPase activity assay sample. The phosphate standards and blank control for colorimetric detection were prepared according to the manufacturer’s instructions of the ATPase/GTPase activity assay kit. An aliquot of 30 μl reaction mix (made with 20 μl assay buffer plus 10 μl 4 mM ATP solution) was added into each ATPase activity assay sample. After incubation at room temperature for 30 min, 200 μl reagent was added to each sample to terminate the enzyme reaction, and all samples were incubated for an additional 20 min. At this step, the transparent reaction mix suddenly changed to a fine green endpoint colour. The colour intensity was measured on a microtitre plate reader at 620 nm. All the assays were repeated five times. The ATPase activity of *Nl*Gr7 was determined by using the mean value of the samples according to the linear regression of standards.

### Polysaccharide (insoluble)-binding study

To identify the specific carbohydrate-binding conserved aromatic amino acid residues in the *Nl*Gr7 sequence, *Nl*Gr7 was aligned with known carbohydrate-binding homologs (**Duan et al., 2016**) to create multiple sequence alignments by using ClustalW (http://www.genome.jp/tools-bin/clustalw). Furthermore, *Nl*Gr7 binding to various insoluble polysaccharides was determined as described by Duan et al (2016). The polysaccharides that were tested were chitin, agarose, Sephadex G-100, raw cassava starch (tapioca starch), alpha-cellulose, and xylan from corn cobs.

### Tastant-binding assay

To study the tastant-binding determination with *Nl*Gr7, the tastant-binding assay with *Nl*Gr7 was performed in 96-well microtitre plates. Microtitre plates were coated with purified *Nl*Gr7 protein (100 ng per well, 100 µl) and incubated overnight (15-16 h, at 4°C). The unbound protein was washed off (flicking and flapping manner) thrice with Tris buffer (25 mM Tris, pH 8.0, 10 mM NaCl). Then, the unbound sites were blocked with 3% bovine albumin serum in Tris buffer and were incubated for 2 h at 100 rpm at room temperature. An additional 100 μl tastants (50, 25, and 12.5 mM) were added into each well and were incubated for 2 h at room temperature. Then, free tastants were washed off thrice with Tris buffer containing 0.05% Tween-20. Subsequently, the wells were incubated with 100 μl biotinylated *Nl*Gr7 (100 ng), followed by incubation with streptavidin-HRP (1: 2000, 1 h, room temperature). Finally, the colour was developed. The bound enzyme activity was measured using 100 μl TMB substrate solution (TransGen Biotech Co., LTD, China) containing 0.06% of 30% v/v hydrogen peroxide (H_2_O_2_) in each well. Due to the enzyme activity, a gradual increase in brilliant blue colour intensity with the increasing concentration of tastants was observed. After 20 min, the reaction was stopped by adding 100 μl sulfuric acid (0.5 N) to each well. In this step, the brilliant blue colour suddenly changed to a fine yellow endpoint colour. The absorbance was measured at 450 nm using an ELISA plate reader.

## Results

### Characterization of *Nl*Gr7

The truncated cDNA of *NlGr*7 consisted of 730 nucleotide bases coding for 243 amino acids with a predicted molecular mass of 28.70 kDa (**Fig. S1**). The cDNA clone was designated *Nl*Gr7 (**Kui et al., 2018**). The estimated pI of the predicted protein *Nl*Gr7 was found to be 9.19. There were eleven polar and nine non-polar amino acid residues. The instability index, as computed by the ExPASy ProtParam tool, was 36.27, which classified the protein as a stable protein (**Fig. S1**). Bioinformatics analysis using the TOPCONS tool predicted the absence of a secretory signal sequence and transmembrane helices in *Nl*Gr7 (**Fig. S2**). *Nl*Gr7 showed a 20.69-22.62% variation in sequence identity with the known Grs sequence of insect pests (**Table S1**). Phylogenetic analysis of *Nl*Gr7 revealed the degree of relationship with respect to genes from other insects (beetle, flies, mosquitos, moths, and butterflies) (**Fig. 1**). However, this study clearly classified *Nl*Gr7 and other insect taxa into two large clades. We employed the polymerase chain reaction to amplify 730 bp *NlGr*7 amplicons. We then cloned, expressed, and purified the amplified amplicons (data not shown). The apparent molecular weight of our protein of interest showed a mobility shift of ∼55.428 kDa on SDS-PAGE due to a high (9.19) isoelectric point (pI) and the presence of hydrophobic amino acids in the *Nl*Gr7 sequence. Furthermore, this purified ∼55.428 kDa protein was confirmed to be *Nl*Gr7 (**Fig. 2A**), which is a sugar transporter of the BPH, based on mass-spectrometry (LCMS/MS-Q-TOF) (**Fig. S3**).

**Figure 1.**
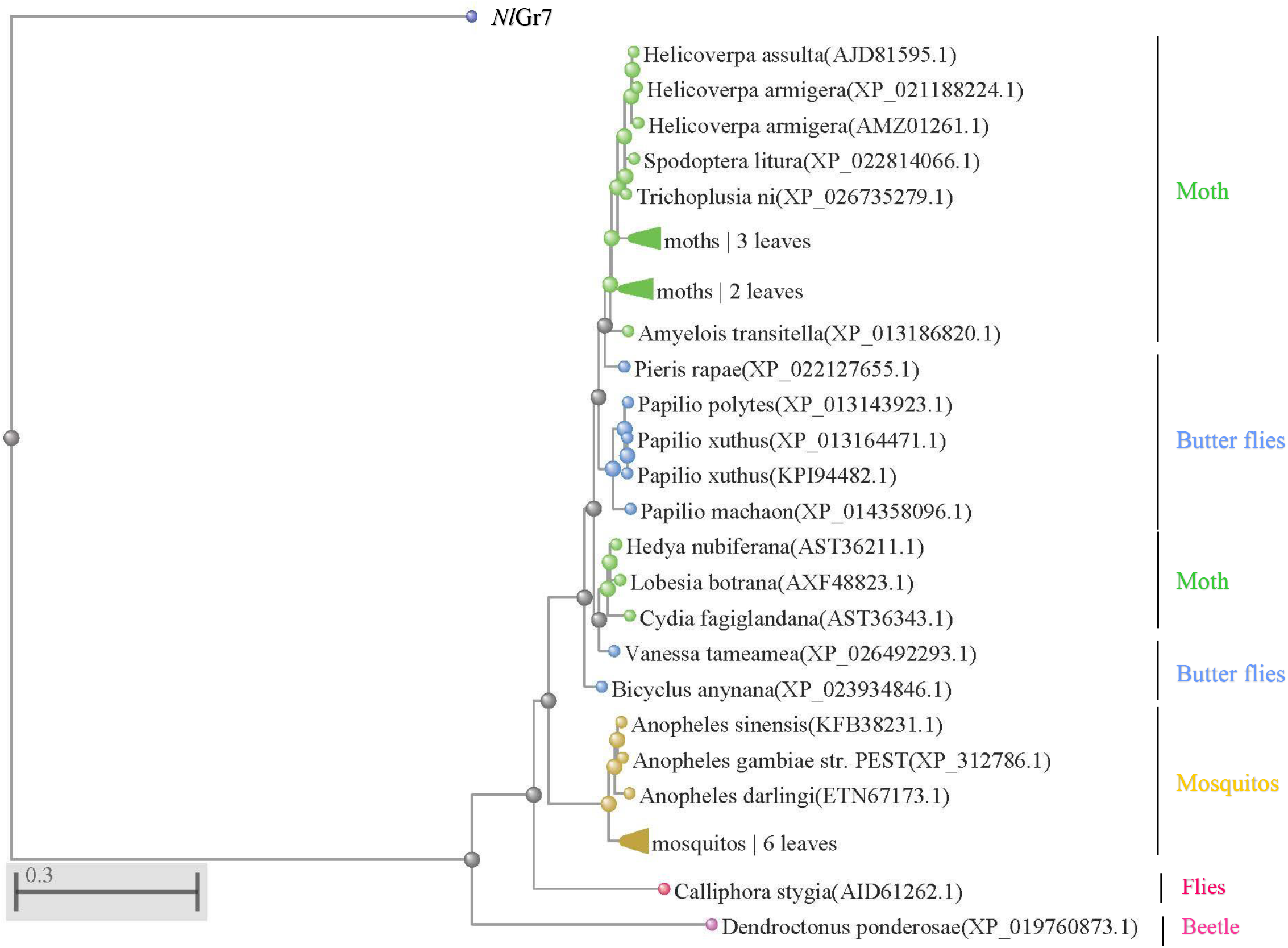
Phylogenetic tree showing the relation between orthologues of gustatory receptor reported from different organisms.

**Figure 2.**
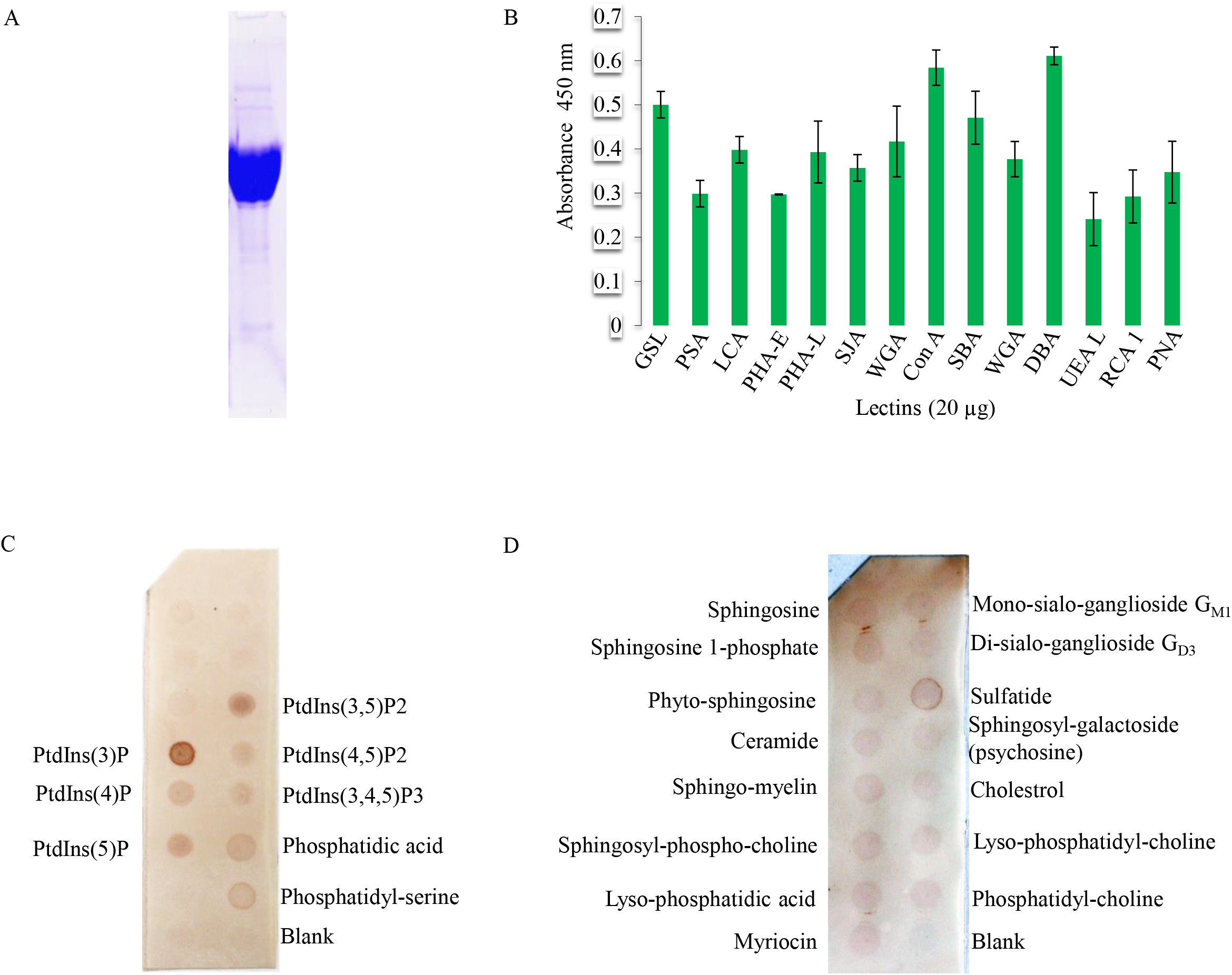
*Nl*Gr7 recombinant protein and *Nl*Gr7-ligands overlay assays. (A) Purified *Nl*Gr7, (B) *Nl*Gr7-lectin overlay assay shows that *Nl*Gr7 binds to several lectins. Protein-lipid overlay assay shows that *Nl*Gr7 binds to several anionic lipids. (C) *Nl*Gr7 reveals interaction to phospholipids, and (D) interaction to sphingolipids.

### Lectin affinity for *Nl*Gr7

To study the affinity of lectins, which are carbohydrate-binding proteins with *Nl*Gr7, a total of fourteen lectins, griffonia (Bandeiraea) simplicifolia lectin (GSL), Pisum sativum agglutinin (PSA), Lens culinaris agglutinin (LCA), Phaseolus vulgaris erythroagglutinin (PHA-E), Phaseolus vulgaris leucoagglutinin (PHA-L), Sophora japonica agglutinin (SJA), wheat germ agglutinin (WGA), concanavalin A (ConA), soybean agglutinin (SBA), wheat germ agglutinin (WGA), Dolichos biflorus agglutinin (DBA), Ulex europaeus agglutinin 1 (UEA1), Ricinus communis agglutinin 1 (RCA1, Ricin), and peanut agglutinin (PNA), were demonstrated to have affinity for *Nl*Gr7. DBA revealed the highest affinity, while UEA1 showed the lowest affinity for *Nl*Gr7 (**Fig. 2B**). The standard deviation (SD) of a set of absorbance values ranged from ±0.001 to ± 0.08 (**Fig. 2B**). These lectins may combine with *Nl*Gr7 and likely play numerous roles in biological recognition phenomena involving cells, carbohydrates, and proteins.

### Lipid affinity for *Nl*Gr7

We screened the arrays of 30 lipids by using the protein-lipid overlay (PLO) assay. This lipid screen showed that recombinant *Nl*Gr7 binds to several anionic phospholipids, including PtdIns(3)P, PtdIns(4)P, PtdIns(5)P, PtdIns(3,5)P2, PtdIns(4,5)P2, PtdIns(3,4,5)P3, phosphatidic acid (PtdOH), phosphatidyl**-**serine (PtdSer) (**Fig. 2C**) and a single strong signal of sphingolipid sulfatide (sulfogalactosylceramide; GalCerI3-sulfate) (**Fig. 2D**).

### Metals affinity to *Nl*Gr7

The metal-binding capability of *Nl*Gr7 was verified by a gel mobility shift assay. The purified *Nl*Gr7 mixed with different concentrations of CuSO_4_ and CaCl_2_ was resolved on SDS-PAGE. Compared with the mobility of *Nl*Gr7 in the presence of 1.0 mM EDTA, the mobility of *Nl*Gr7 was slowed by the addition of 0.015 to 1.0 mM CuSO_4_. The migration of *Nl*Gr7 appeared to slow when the concentration of CuSO_4_ was high (**Fig. 3A**), which suggested that *Nl*Gr7 has the ability to bind Cu^2+^. Furthermore, the copper-binding site, as predicted by the RaptorX-binding server (http://raptorx.uchicago.edu/BindingSite),was interesting for the observation of glutamine (Q)-64, asparagine (N)-117, lysine (K)-218, glutamine (Q)-222, threonine (T)-225, tyrosine (Y)-226, and isoleucine (I)-229 in the *Nl*Gr7 domain (**Fig. 3B**). The mobility of *Nl*Gr7 shifted with the addition of CaCl_2_ in comparison with the mobility of *Nl*Gr7 in the presence of 1.0 mM EDTA (**Fig. S4**). On SDS-PAGE, the calcium-*Nl*Gr7 complex was stable in the presence of a low (0.015 and 0.05 mM) CaCl_2_ concentration. At the same time, the calcium-*Nl*Gr7 complex was reduced in the presence of increasing (0.15, 0.5, and 1.0 mM) concentrations of CaCl_2_ (**Fig. S4**). This reduced intensity of the calcium-*Nl*Gr7 complex on SDS-PAGE was due to the precipitation of the *Nl*Gr7 protein in the presence of a high concentration of calcium chloride under incubation for 30 min at 25°C.

**Figure 3.**
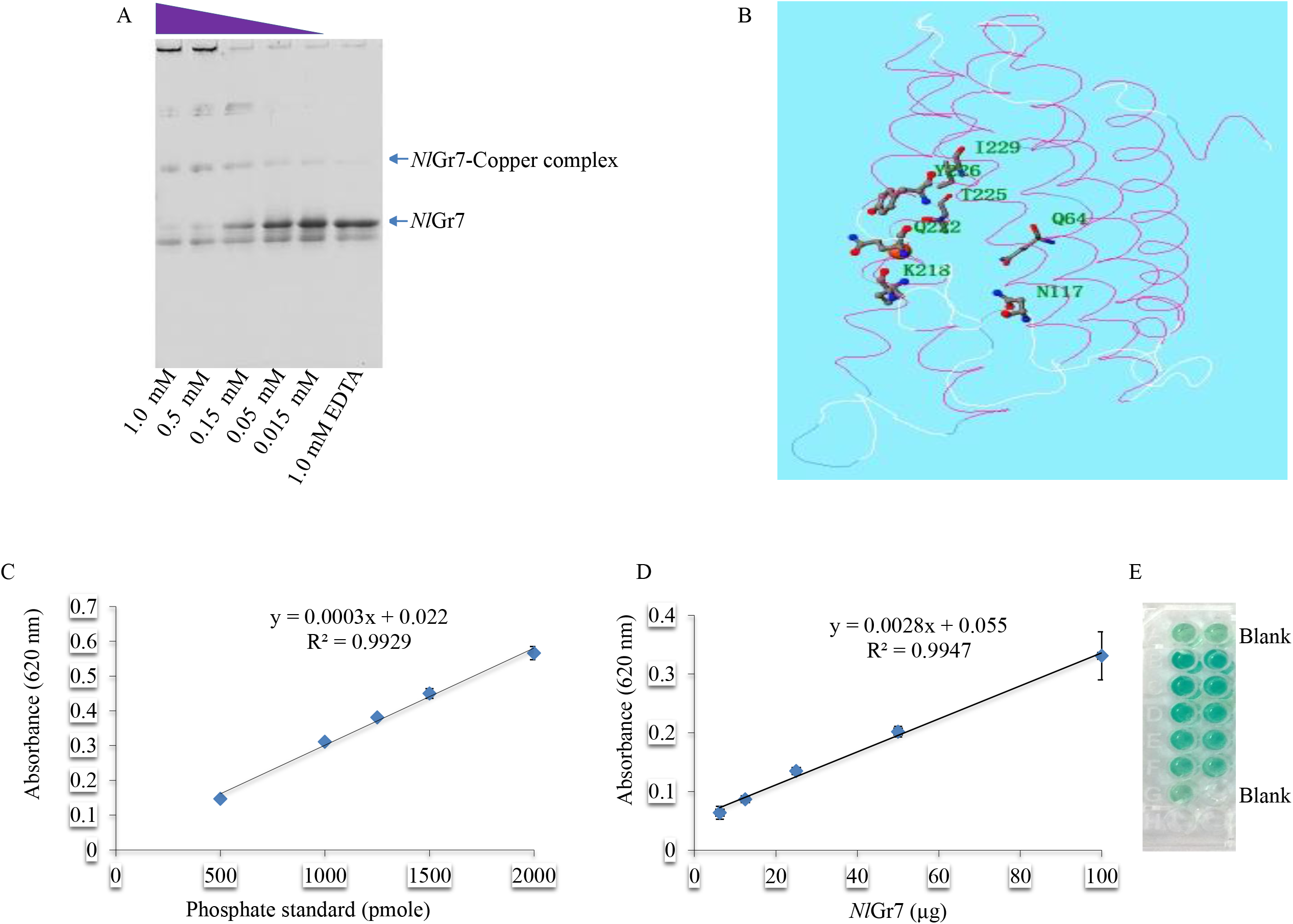
Overview of the *Nl*Gr7-copper complex and ATPase/GTPase activity assay for *Nl*Gr7. (A) The gel mobility shifts of the *Nl*Gr7-copper complex showed a varied range of copper. (B) Predicted binding amino acid residues Q64, N117, K218, Q222, T225, Y226, and I229 anchor copper within the *Nl*Gr7. (C) Standard curve prepared according to the ATPase/GTPase activity kit instructions. (D) The *Nl*Gr7 (6.25, 12.5, 25, 50, or 100 µg) determined ATP. (E) This ATPase/GTPase activity assay is a colorimetric test that shows the amount of free inorganic phosphate (Pi). Controls that included no *Nl*Gr7 were measured to observe for spontaneous ATP hydrolysis of ATP, and their optical density (OD) values were deducted from the samples’ values.

### Hydrolysis of the *Nl*Gr7-bound ATP

To understand how the predicted active site of *Nl*Gr7 might relate to its biological function, we investigated its biochemical activities *in vitro*. A total of 4 phosphorylated sites (serine (S)-54, tyrosine (Y)-104, tyrosine (Y)-105, and threonine (T)-201) and their corresponding catalytic protein kinases were predicted from the *Nl*Gr7 amino acid (1-243) sequence (http://kinasePhos.mbc.nctu.edu.tw/, **Fig. S5**). Furthermore, the developed phosphate standard curve showed that phosphate could be detected at a minimum of 500 pmol and maximum of 2000 pmol. The R^2^ value was 0.992, as shown in the figure (**Fig. 3C, Table S2A**). Examination of the ATPase activity assay of bound phosphate in *Nl*Gr7 showed that 217.68-1125.85 pmol of phosphate was liberated (**Fig. 3D, Table S2B**), and its respective stable dark green colour with free phosphate liberated by the enzyme in a colorimetric product is shown in the figure (**Fig. 3E**). The R^2^ value was 0.994, as shown in the figure (**Fig. 3D**). These data are consistent with the finding that *Nl*Gr7 displays its ATPase activity *in vitro*. We postulated that phosphate binding plays an important biochemical function.

### *Nl*Gr7 exhibited polysaccharide (insoluble) specificity

The purified *Nl*Gr7 showed a binding affinity for six insoluble polysaccharides. *Nl*Gr7 showed the highest affinity for tapioca starch, while *Nl*Gr7 showed the lowest affinity for Sephadex G-100 (**Fig. 4A**). Furthermore, seven carbohydrate-binding homologs sequences (**Duan et al., 2016**), belonging to accession numbers ADR64668-2, AFN57700-2, CBM_C5614- 1_, ADR64664, ADR64668-1, AFN57700-1, and CAJ19146, were aligned with *Nl*Gr7 to observe conserved aromatic amino acids, which were accepted to play an essential role in identifying and binding to polysaccharides. Two (Tryptophan (W)-59, and W-128) aromatic amino acids were identified as being completely conserved in all the aligned-sequences (numbered according to amino acids in *Nl*Gr7) (**Fig. 4B**, Red box). At the same time, phenylalanine (F)-91, tyrosine (Y)-104, F-108, and F-111 were observed to be partially conserved aromatic amino acids in all the aligned sequences (**Fig. 4B**, Pink box). A phylogenetic tree revealed that *Nl*Gr7 showed the least similarity with other known carbohydrate-binding homolog sequences (**Fig. 4C**).

**Figure 4.**
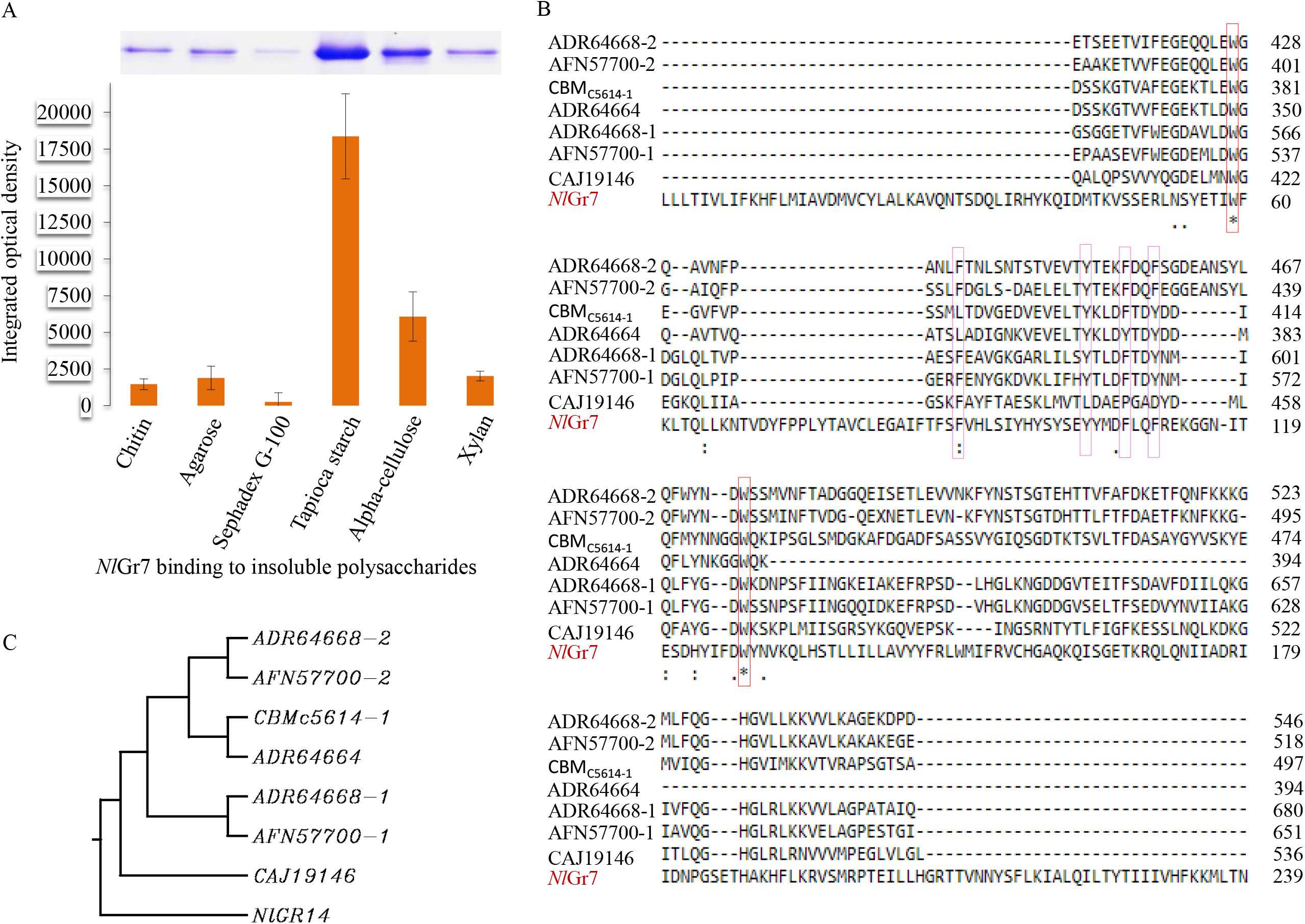
Characterization of *Nl*Gr7 as a novel carbohydrate protein. (A) Binding of *Nl*Gr7 to insoluble polysaccharides. Thirty micrograms of purified *Nl*Gr7 was incubated with 200 μl 4% (wt/vol) insoluble polysaccharide including chitin, agarose, Sephadex G-100, raw cassava starch (tapioca starch), alpha-cellulose, or xylan from corn cobs. The same amount of protein used in the binding assay but without polysaccharide was included as a control (CK). (B) Multiple sequence alignment of *Nl*Gr7 and selected carbohydrate-binding homologs. The sequence similarity and identity are shown by dots and asterisks, respectively. Entirely conserved aromatic (tryptophan (W)-59, and W-128) amino acids are red, and partly conserved aromatic (phenylalanine (F)-91, tyrosine (Y)-104, F-108, and F-111)amino acids are pink. (C) Phylogenetic tree showing the relation between orthologues of the carbohydrate protein that was reported previously.

### *Nl*Gr7 showed an interaction with various tastants

Twelve tastants (D-(+) glucose, maltose, sucrose, D-(+) galactose, D-xylose, trehalose, D-(-) ribose, D-(-) melezitose, maltotriose, D-sorbitol, D-cellobiose, and myoinositol) were individually analysed, and all showed binding affinity for *Nl*Gr7 (**Fig. 5**). The binding affinities of the tastant preparations at each of the 3 concentrations of 50, 25, and 12.5 mM are depicted in Figure 5. Among the tastants, 50 mM D-cellobiose showed the highest (absorbance 0.352) binding affinity for *Nl*Gr7 (**Fig. 5K**), while 50 mM sorbitol showed the lowest (absorbance 0.065) binding affinity for *Nl*Gr7 (**Fig. 5J**). The standard deviation (SD) of a set of absorbance values ranged from ±0.0005 to ± 0.042 (**Fig. 5**).

**Figure 5.**
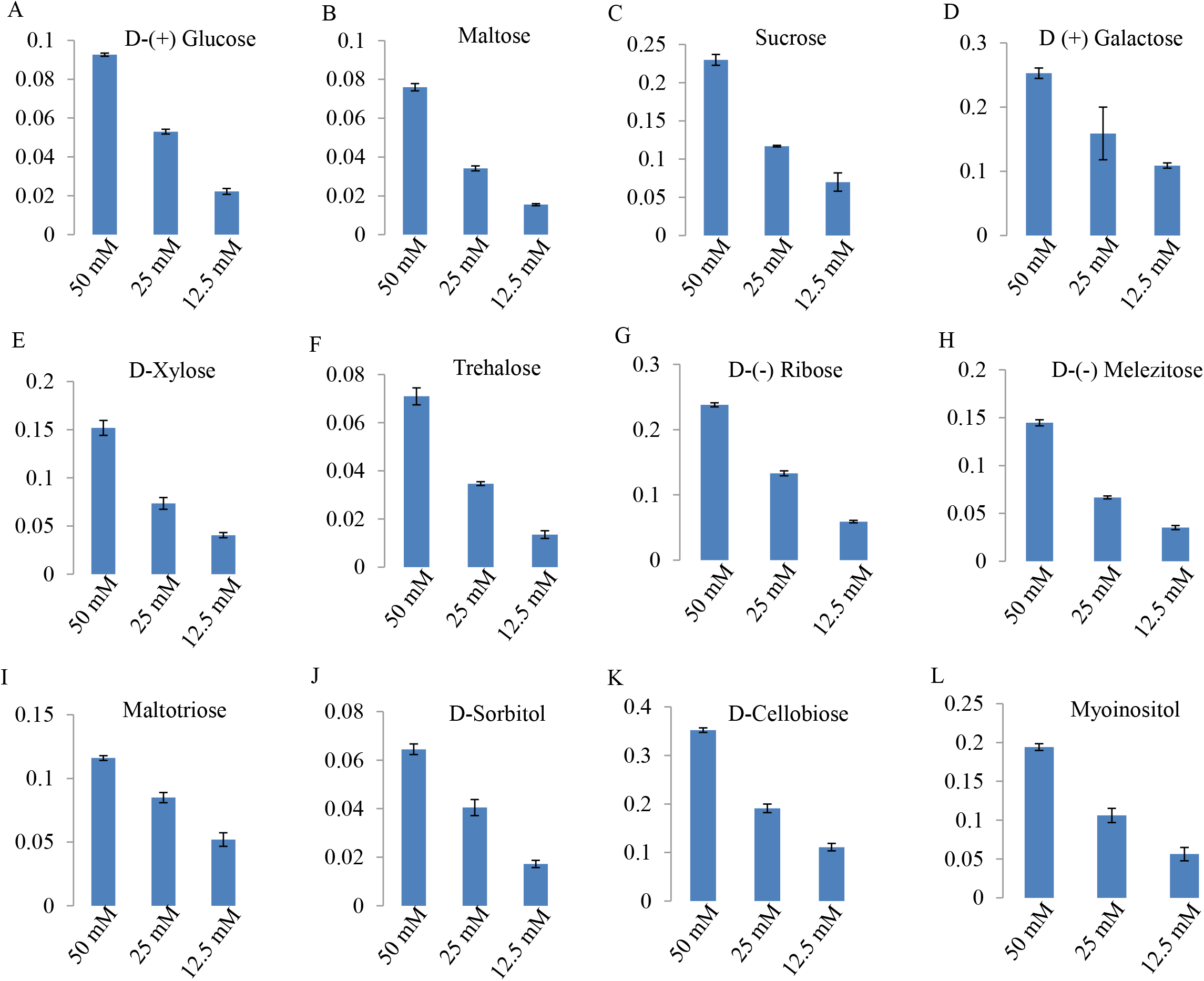
Tastants screening assay. The binding of *Nl*Gr7 to different concentrations (50, 25, and 12.5 mM) of tastants. (A) D-(+) glucose, (B) maltose, (C) sucrose, (D) D(+) galactose, (E) D-xylose, (F) trehalose, (G) D-(-) ribose, (H) D-(-) melezitose, (I) maltotriose, (J) D-sorbitol, (K) D-cellobiose, and (L) myoinositol.

## Discussion

Here, we systematically characterized the functional properties of *NI*Gr7 using the macromolecular (*in vitro*) approach. Our study begins to overcome challenges in studying this highly divergent superfamily of insect Gr proteins and provides a systematic overview of ligand (metals, carbohydrates, lipids, lectins, and tastants) detection and ATP binding by this sugar receptor. The majority of gustatory receptor ligands in *Drosophila* have been studied by using electrophysiology recordings and/or behavioural investigations. However, there is, to some extent, a gap in the accurate interpretations of the electro-physiological data and behaviours that are recorded between wild-type and Gr mutant flies (**Miyamoto et al., 2013**). The lectin-binding proteins of the rice brown planthopper are still undefined. Primary information from the BPH gustatory receptor and the lectin(s)-binding study reveals that the structure that interacts with the lectin(s) consists of carbohydrate moieties (**Lis and Sharon, 1986**). A variety of carbohydrate specificity-lectin(s) were selected for study (**Fig. 2B**). In assays, the *Nl*Gr7 protein interacted with a variety of sugar-specific lectins, which offers a complex oligosaccharide structure. Similar facts were found earlier for the human parotid salivary gland (**Rohringer and Holden, 1985**). The role that lectin-binding with *Nl*Gr7 (protein) plays in the developmental stages of the BPH has yet to be determined.

Although phospholipids (PLs) are only a minor nutrient compared to starch and protein, they may have both nutritional and functional significance. Phosphatidylcholine (PC), phosphatidylethanolamine (PE), phosphatidylinositol (PI) and their lyso forms are the major PLs in rice. Two PLs, PtdIns4P (phosphatidylinositol monophosphate) and PtdIns(4,5)P2 (phosphatidylinositol bisphosphate), have been reported as key intermediate sources of second messengers, and they directly affect the signal pathways in many animal cells and plants cells (**Chen et al., 1991**). In this study, *Nl*Gr7 binds to various lipids with different affinities and specificities (**Fig. 2C, D**), which could be the result of non-specific electrostatic interactions. The noticed affinities of *Nl*Gr7 for the lipids are specifically known as PH (pleckstrin homology) lipid-binding domains (**Stahelin, 2009**). However, various examples in prior studies show that low-affinity interactions have a distinct and influential impact on biological roles (**Becker et al., 1996**; **Irwin and Tan, 2014**; **Wilson, 2003**). Our rationale, which can be noticed in *Nl*Gr7-lipid interactions and in the lipids with their reported biological functions, reveals that lipid binding targets the chaperone proteins to specific cellular locations, where, in addition to participating in transmembrane targeting and protein folding, they also affect the physiological pathway of the membrane. Hence, the specific binding of lipids suggests that lipid binding has been a conserved function of *Nl*Gr7 since its appearance (**Fig. 2C, D**).

Copper is an important micronutrient that plays an essential role in plant growth and development (**Roy, 1931**); little is known about copper in insects. Insects often consume heavy metal ions from their niche. Previous reports revealed that copper is related to both the developmental stage of the organism, when animals undergo the transition from larval to adult blood types (**Durstewitz and Terwilliger, 1997**; **Ye et al., 2015**; **Li et al., 2016**), and the moulting cycle, when large fluctuations in protein synthesis take place (**Ye et al., 2015**; **Li et al., 2016**). In the present study, *Nl*Gr7 showed an interaction with copper. Therefore, taking this observation further, we speculate that copper may also play a role in the developmental stage (egg to adult) and moulting cycle of BPH. However, this has yet to be verified.

It has been noted that effectors from herbivores, which can bind to Ca^2+^, are involved in suppressing the plant’s defence (**Atamian et al., 2013**) and contracting forisomes (**Will et al., 2007**). However, the physiological mechanism of effector-mediated regulation of plant defences remains widely unexplained (**Consales et al., 2012**). The structure of the transduction pathway in gustation is unclear. The signalling pathways, including cGMP (**Amakawa et al., 1990**), inositol 1,4,5-triphosphate (IP_3_) (**Koganezawa and Shimada, 2002**), and the sugar-receptor protein-gated channel (**Murakami and Kijima, 2000**), have been studied in few insects. Furthermore, GPCRs activate a temporal rise in intracellular Ca^2+^ because inositol 1,4,5-triphosphate activates delivery of Ca^2+^ from intracellular storage (**Torfs et al., 2002**). Therefore, we speculate that the GPCRs-IP_3_-Ca^2+^-mediated network may impact the regulation of ligand-mediated calcium flux in the BPH physiological mechanism, but this has yet to be elucidated.

Polysaccharides display a wide range of solubility in water. However, some are water insoluble, e.g., chitin, agarose, tapioca starch, alpha cellulose, and xylan. Cellulose and chitin polysaccharides play structural roles in plants, fungi, and insects. The interactions between proteins and cellulose (**Georgelis et al., 2012**; **Boraston et al., 2004**; **Boraston, 2004**), as well as between human chitin-binding proteins (YKL-39 and −40) (**Schimpl et al., 2012**), have been studied extensively. However, the physiological role of these proteins remains poorly understood. Our results revealed a new carbohydrate-binding protein, *Nl*Gr7, which targets different insoluble polysaccharides that adopt varied conformations, including chitin, agarose, tapioca starch, alpha cellulose, and xylan. This study reveals that *Nl*Gr7 may possess higher plasticity to accommodate varied ligand conformations. This varied binding specificity remains unexplored without a structural study of the *Nl*Gr7-ligand complex, and it has yet to be a priority in future studies.

To study *Nl*Gr7 as a sugar Gr, we tested three different concentrations of twelve sugars to reveal the receptor’s ligands. The results revealed that all 12 sugars interacted with *Nl*Gr7 (**Fig. 5**). A previous report demonstrated that galactose, xylose, glucose, and trehalose can regulate intracellular Ca^2+^ levels in insect cell lines (**Chen et al., 2019**; **Xu et al., 2012**). We speculate that expression of *Nl*Gr7 within the BPH may regulate the expression, localization, and/or function of the endogenous taste receptors of the BPH, thus leading to an increase or decrease in the responses to tastants.

## Conclusions

Currently, there is insufficient knowledge about the function of gustatory (sugar) receptors within the insect. In a previous study in *Drosophila*, Gr43a showed upregulation during the feeding experiences of hungry flies and downregulation during feeding experiences in satiated flies (**Miyamoto et al., 2012**). In the present study, we observed the functional versatility of *Nl*Gr7. We assume that *Nl*Gr7 binding with its ligands may likely stimulate the physiological pathways of the BPH for feeding behaviour and adaptation. These observations extended our knowledge of the Grs (proteins) mechanism in insect and pest control.

## Author contributions

WZ and AO conceived and designed the experiments. AO performed the experiments and analyzed the data. AO and WZ wrote the manuscript. Both authors have read and approved the final manuscript.

## Acknowledgements

We thank Longgyu Yuan for their assistance with insect maintaining in green house at State Key Laboratory of Biocontrol, SunYat**-**sen University, China.

## Supplementary materials

**Figure S1**. The computation of various physical and chemical parameters of *Nl*Gr7.

**Figure S2**. Consensus prediction of membrane protein topology using the TOPCONS server indicated no trans-membrane helices (blue in the graph) in the predicted amino acid sequence of *Nl*Gr7. TOPCONS predicted the topology of *Nl*Gr7 from five different topology prediction algorithms: OCTOPUS, Philius, PolyPhobius, SCAMPI (multiple sequence mode), and SPOCTOPUS. The output of these five algorithms was used as input for the TOPCONS Hidden Markov Model (HMM) (shown in maroon), which provided a consensus prediction for the protein together with a reliability score based on the agreement of the included methods across the sequence. In addition, ZPRED was used to predict the Z-coordinate (i.e., the distance to the membrane centre) of each amino acid, and the G-scale was used to predict the free energy of the membrane insertion for a window of 19 amino acids that were centred around each position in the sequence.

**Figure S3**. LCMS/MS-Q-TOF-based identification of the *Nl*Gr7. (A) Parameters used in MASCOT search, (B and C) The green area is a significant threshold. This area of the score over the results of the identification of the mascot positive result**s**, (D and E) decoy search summary, and (F) cross-examination of the LCMS/MS-QTOF-generated peptide sequence of the sugar transporter sequence blast in the NCBI protein database.

**Figure S4**. An electrophoretic mobility shift assay (EMSA) for the calcium-binding assay.

**Figure S5**. Computationally predicted catalytic kinase-specific phosphorylation sites within the *Nl*Gr7 sequence using the KinasePhos tool.

**Table S1**. NlGr7 protein sequence identity with known chemoreceptors of insects.

**Table S2**. ATPase activity assay with NlGr7. Hydrolysis of the N*l*Gr7-bound ATP. (A) Standard curve of phosphate, (B) the concentration of released phosphate from the N*l*Gr7 bound ATP. The released concentration of phosphate from the samples per well was calculated using the phosphate standard curve value 500 pmol.

